# Gadusol is a maternally provided sunscreen that protects fish embryos from DNA damage

**DOI:** 10.1101/2023.01.30.526370

**Authors:** Marlen C. Rice, Jordan H. Little, Dale L. Forrister, Julane Machado, Nathan L. Clark, James A. Gagnon

**Affiliations:** School of Biological Sciences, University of Utah, SLC, UT 84112, USA; Department of Human Genetics, University of Utah, SLC, UT 84112, USA; Henry Eyring Center for Cell & Genome Science, University of Utah, Salt Lake City, UT 84112, USA

## Abstract

Ultraviolet radiation (UVR) and its deleterious effects on living cells selects for UVR-protective mechanisms. Organisms across the tree of life evolved a variety of natural sunscreens to prevent UVR-induced cellular damage and stress. However, in vertebrates, only melanin is known to act as a sunscreen. Here we demonstrate that gadusol, a transparent compound discovered over 40 years ago in fish eggs, is a maternally provided sunscreen required for survival of embryonic and larval zebrafish exposed to UVR. Mutating an enzyme involved in gadusol biosynthesis increases the formation of cyclobutane pyrimidine dimers, a hallmark of UVB-induced DNA damage. Compared to the contributions of melanin and the chorion, gadusol is the primary sunscreening mechanism in embryonic and larval fish. The gadusol biosynthetic pathway is retained in the vast majority of teleost genomes but is repeatedly lost in species whose young are no longer exposed to UVR. Our data demonstrate that gadusol is a maternally provided sunscreen that is critical for early-life survival in the most species-rich branch of the vertebrate phylogeny.

## Introduction

Most life on earth relies on photosynthetic food webs for their energy source, which can result in extensive exposure to ultraviolet radiation (UVR)^1^. UVR, especially UVB (280-320nm), can damage proteins and DNA, leading to errors during DNA repair and replication. Excessive UVR induces cellular death. Aquatic organisms risk UV exposure because biologically harmful levels of UVB can penetrate >10 m in clear water^2^. Organisms in diverse habitats adapt to avoid, ameliorate, or protect against the effects of UVR. Some of these adaptations include sun avoidance behaviors (e.g., nocturnal lifestyle) and DNA repair machinery (e.g., nucleotide excision repair)^3^. However, since sunlit habitats can have significant nutritive advantages over dark environments and because no repair pathway is completely efficient, many organisms employ sunscreens to avoid UVR damage from occurring in the first place^4^.

Sunscreens absorb UV photons before they penetrate vulnerable cells and dissipate this absorbed energy as less harmful heat. A wide variety of sunscreens are used by living organisms, including flavonoids in plants, scytonemin in cyanobacteria, and melanin in numerous organisms including vertebrates^4^. In fish and other aquatic vertebrates, melanin is produced in melanophores (homologous to melanocytes in mammals), which differentiate from embryonic neural crest cells and migrate to cover aspects of the brain and body^5^. Recently, an internal melanophore umbrella was shown to protect the hematopoietic stem and progenitor cell niche in developing zebrafish from UVR^6^. However, since melanophores emerge late in embryonic development, they cannot protect early stages when the embryo is most sensitive^7^, and some controversies remain about melanin’s sunscreening role in fish^8–10^. Thus, the mechanisms that may protect the initial phases of development in externally fertilized vertebrate embryos (e.g., the vast majority of fish) remain mysterious. Apart from melanin, no endogenously produced sunscreen has been documented in vertebrates.

Mycosporine-like amino acids (MAAs) are a class of sunscreening compounds produced by numerous algae and microbes^8^. Experiments indicate that depletion of MAAs causes UVR sensitivity in cyanobacteria^11^and sea urchins^12,13^. The eggs of many fish species also contain large quantities of an MAA-related UVR-absorbent compound called gadusol (first discovered in eggs of the cod *Gadus morhua*)^14,15^. Although the existence of gadusol in fish eggs and embryos was discovered decades ago, its role as a sunscreen remains untested^16^. Gadusol in fish was originally thought to come from dietary sources^1,2^. However, a two-gene cassette was recently discovered in many vertebrate genomes that enables the production of gadusol from sedoheptulose-7-phosphate (an intermediate in the pentose phosphate pathway)^17^. Yeast engineered to express the zebrafish biosynthetic pathway produced gadusol, which provided protection against UVR in yeast^17^. With an understanding of the teleost genetic architecture underlying gadusol production, and a genome-editing toolkit for zebrafish, it is now possible to test the role of gadusol as a sunscreen *in vivo*.

Here, we test the role of gadusol in UVR protection of fish embryos and larvae by generating a gadusol-deficient mutant zebrafish. We determine that gadusol is maternally provided in embryos and provides protection from UVR throughout embryonic and larval development. We find that gadusol is the primary sunscreen during fish development while melanin and other mechanisms provide secondary protection. In a broader evolutionary context, we also find a striking pattern of repeated loss of gadusol production in fish species whose embryos are not exposed to sunlight. Together, our work provides evidence that gadusol is a widely distributed and evolutionarily conserved sunscreen that protects vertebrate embryos in aquatic sun-lit environments.

## Results

### Gadusol is maternally provided and protects embryos and larvae from UVR

To test if gadusol is a sunscreen in vertebrate embryos, we used CRISPR-Cas9 to delete most of exon 2 of zebrafish *eevs*, which encodes the enzyme essential for the first step in gadusol biosynthesis (**Fig. S1**). We chose zebrafish for these experiments because they live and spawn in shallow sunlit waters, they are known to produce gadusol^17^, and they are genetically tractable. Grown in our animal facility, where they are protected from UVR, homozygous *eevs* mutant females and males survived to fertile adulthood like their wild-type peers. Using reciprocal crosses between homozygous mutant adults (*eevs*^-/-^) and wild-type adults (*eevs*^+/+^), we generated heterozygous mutant embryos that lack maternal contribution of gadusol (hereafter referred to as M*eevs*) and heterozygous mutant embryos that retain this maternal contribution (referred to as *eevs*^+/-^) (**Fig. 1A**). Notably, M*eevs* and *eevs*^+/-^ embryos have identical genotypes but either lack or possess maternally provided gadusol, as judged by mass spectrometry (**Fig. 1A**) and UV-spectrophotometry (**Fig. S2**). We generated maternal-zygotic homozygous mutant embryos (referred to as MZ*eevs*) from in-crosses of homozygous mutant parents. Immediately after fertilization, gadusol was nearly absent in MZ*eevs* embryos and indistinguishable from M*eevs* (**Fig. 1B**).

**Figure 1.**
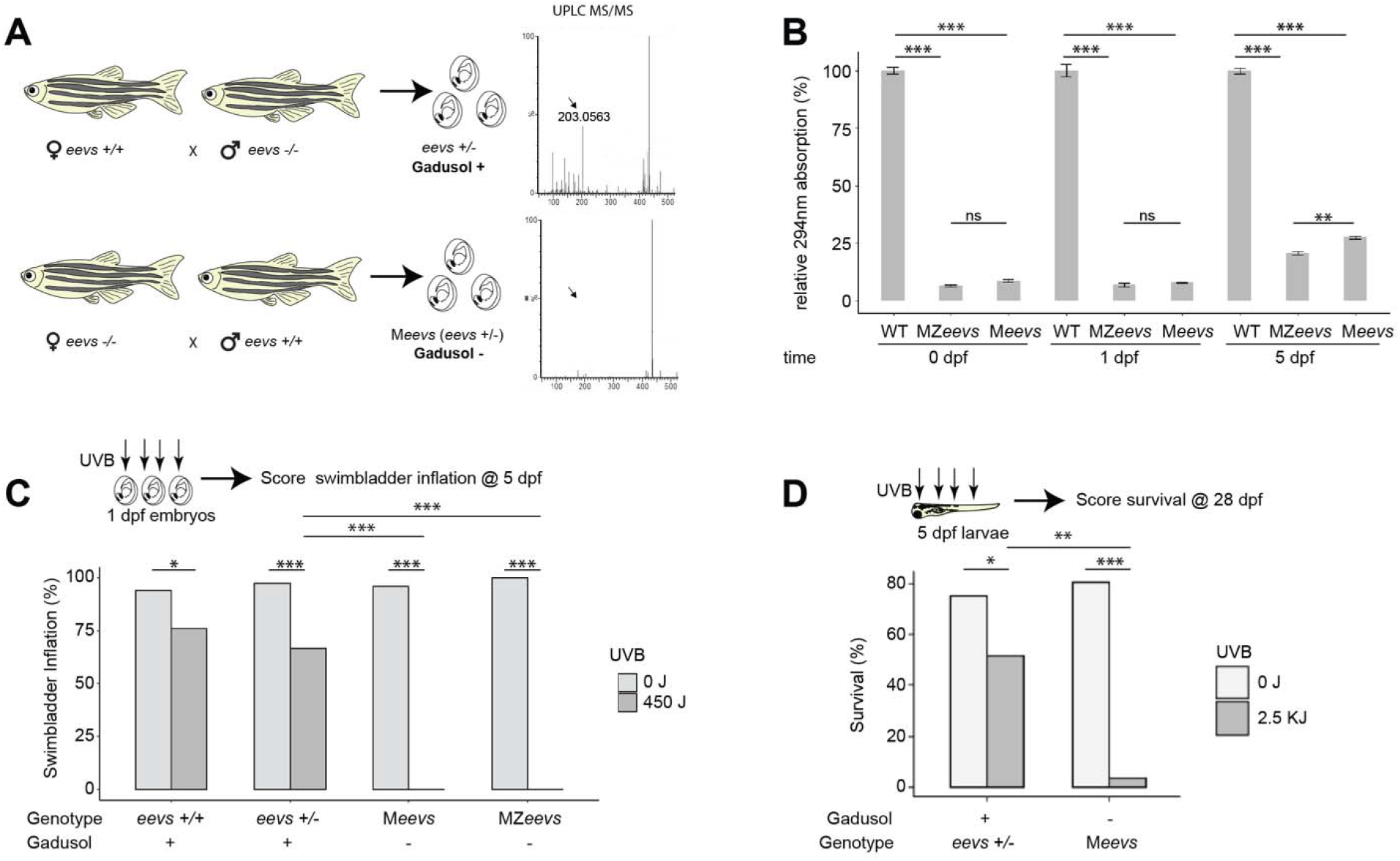
Gadusol is maternally provided and protects zebrafish embryos and larvae from UVB. **A.** Experimental diagram for generating heterozygous mutant *eevs*^+/-^ embryos and larvae with identical genotypes but containing maternal contribution of gadusol (top), or depleted of maternally provided gadusol (bottom). On the right, UPLC mass spectra of 0hpf egg extracts from each genetic cross; arrow indicates gadusol mass. **B.** Absorption values at 296nm from the indicated genotypes at the indicated timepoints. All absorption values normalized to wild type. Error bars indicate standard deviation from biological replicates. **C.** Distribution of swimbladder inflation scored in 5 dpf larvae, with genotypes and gadusol presence indicated, after mock exposure (grey) or UVB exposure (dark grey) at 24 hpf stage. All embryos resulted from crosses between TU and AB strain parents, except the TU in-cross that generated MZ*eevs* embryos. From left to right, n = 50, 50, 75, 75 100, 97, 50, 50; N = 2, 2, 3, 3, 4, 4, 2, 2. **D.** Survival distribution scored at 28 dpf, with genotypes and gadusol presence indicated, after mock exposure (grey) or UVB exposure (dark grey) at 5 dpf. From L-R n = 100, 95, 100, 97, N = 4 for all groups. n = embryos/larvae. N = clutches. statistics: student t test b, Fisher’s Exact t-test c, d, *p<0.05; **p<0.01; ***p<0.0001.

We next asked how long maternally provided gadusol persisted in embryos and larvae. We compared gadusol abundances from whole embryos and larvae with the following genotypes: *eevs*^+/+^ (wild-type), M*eevs*, and MZ*eevs*. We found only a modest increase in gadusol abundance in M*eevs* relative to MZ*eevs* at 5 days post-fertilization (dpf), suggesting that maternally synthesized and deposited gadusol is the source of nearly all gadusol in the developing zebrafish (**Fig. 1B**). This is an example of a maternal effect, where disruption of the *eevs* gene in mothers eliminates gadusol presence in their embryos, regardless of embryo genotype.

To determine if gadusol protects zebrafish embryos against UVB, we developed an assay to deliver precise doses of UVB to embryos and measure the effect on swim bladder inflation at 5 dpf (a hallmark of healthy development essential for survival, **Fig. S3**). We found that 450 joules (J)/m^2^ of UVB (fluence rate: 2.5 W/m^2^, see **Methods**) delivered at 24 hours post-fertilization (hpf) resulted in ~75% swim bladder inflation in wild-type and *eevs*^+/-^ embryos, respectively, but did not result in gross developmental defects (**Fig. S4**). In stark contrast, MZ*eevs* and M*eevs* embryos were extremely vulnerable to the same dose of UVB; all embryos failed to inflate their swim bladders (**Fig. 1C**).

Since zygotic production of gadusol was still minimal at 5 dpf (**Fig. 1B**), we hypothesized that larvae lacking maternal gadusol should be highly sensitive to UVB at this later stage. We repeated UVB dosage curves on 5 dpf larvae and identified 2.5 kJ for a small but significant impact on wild-type larvae survival (**Fig. S5**). We grew UV-exposed and control larvae in our fish facility nursery to 28 dpf, which requires developing animals to forage for food to survive. We found that only 2% of exposed M*eevs* larvae survived, compared to ~50% of controls exposed to the same dose of UVB (**Fig. 1D**). Together, these data demonstrate that maternally provided gadusol provides powerful UVB protection to early embryos and older larvae.

### Gadusol prevents DNA damage and apoptosis

Next, we sought to understand the mechanism by which gadusol protects embryos from UVB. In other species, gadusol and related molecules were hypothesized to function as antioxidants as well as sunscreens^14,17,18^. To test if gadusol serves as an antioxidant in zebrafish embryos, we exposed 24 hpf embryos to hydrogen peroxide to induce oxidative stress. At 5 dpf, gadusol-depleted M*eevs* and control *eevs*^+/-^ embryos had similar responses to increasing doses of oxidative stress, suggesting that gadusol does not function as an antioxidant *in vivo* (**Fig. S6**).

To test if gadusol serves as a sunscreen by absorbing UVB, we measured the production of cyclobutane pyrimidine dimers (CPDs), a signature of UVB-induced DNA damage^19^. If gadusol acts as a sunscreen, then it would absorb UVB photons and shield the underlying DNA from CPD formation. We exposed 24 hpf embryos to UVB and used immunohistochemistry to detect CPDs and quantify fluorescence intensity. Embryos that lacked gadusol had significantly higher levels of CPD formation after UVB exposure compared to controls containing gadusol (**Fig. 2A,B**). CPDs are cytotoxic and at high abundance induce apoptosis. We used immunohistochemistry to detect a fast-acting apoptotic marker (activated caspase-3) in embryos exposed to UVB^20^ (**Fig. 2C, Fig. S7**). We found that embryos lacking gadusol had increased levels of apoptotic nuclei, relative to controls (**Fig. 2C**), supporting a role for gadusol in absorbing UVB and preventing DNA damage.

**Figure 2.**
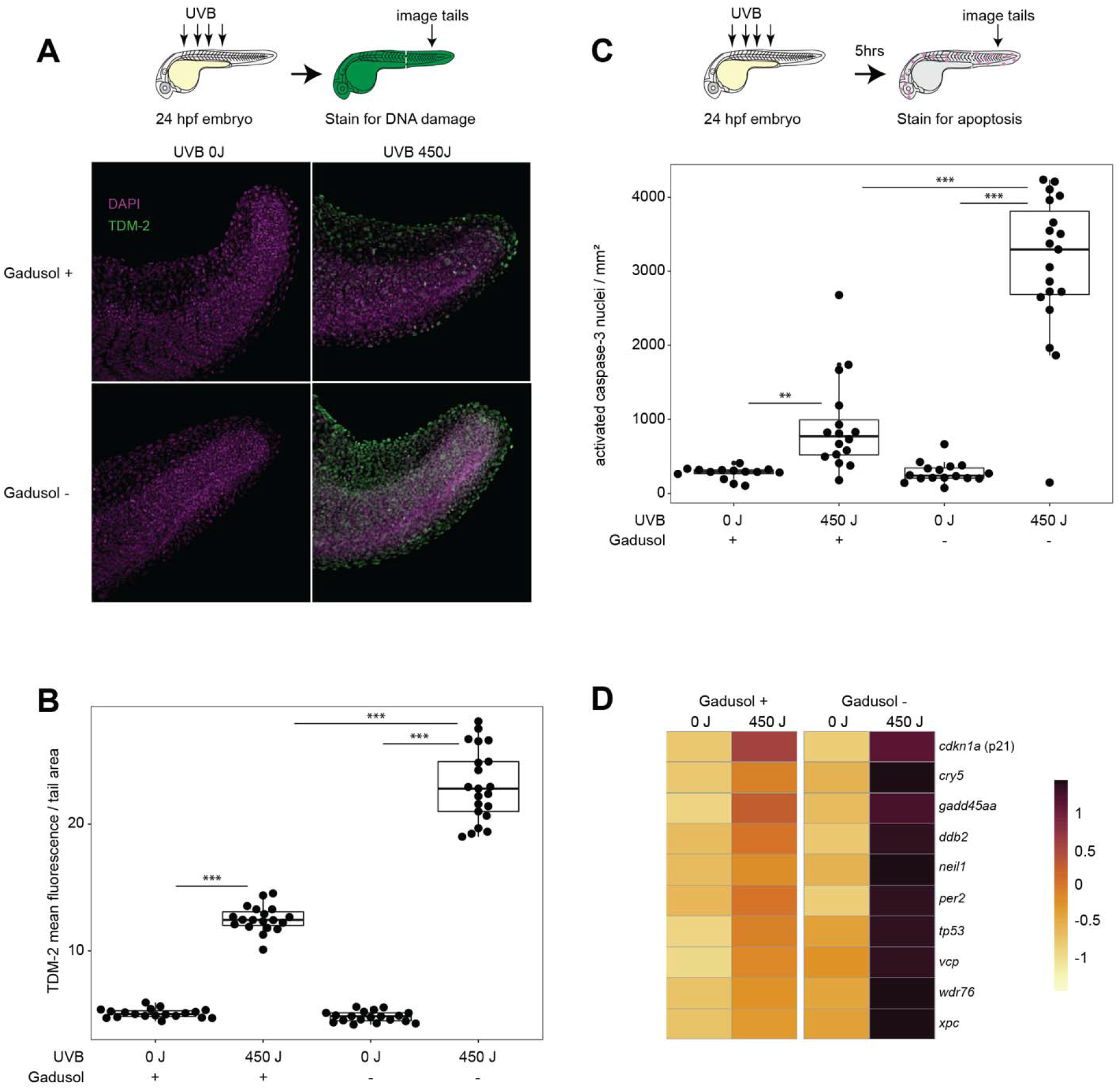
Gadusol functions as a sunscreen preventing DNA damage and apoptosis. **A.** Immunohistochemistry, using an antibody that recognizes CPDs (TDM-2), on 24 hpf immediately after mock or UVB exposure. Representative images shown. **B.** Quantification of CPD labeling normalized to tail area (mm^2^). From left to right, n = 19, 19, 19, 21; N = 2 for all groups. **C.** Quantification of immunohistochemistry, using an antibody that recognizes activated caspase-3. n = 14, 16, 16, 20. N = 2 for all groups. **D.** Significant upregulation of select UVR response and DNA damage GO term-associated genes measured from the indicated conditions and genotypes using RNAseq on 24 hpf embryos after mock exposure or UVB exposure. RNA was collected 5 hours post mock or UVB exposure. Gene expression is scaled by rows. Significance determined via Fishenricher^42^. Student’s T-test P*<0.05; P**<0.01; P***<0.0001. n = number of embryos. N = number of clutches.

To characterize transcriptional responses to UVR in the absence of gadusol, we performed RNAseq comparing gadusol-depleted M*eevs* and wild-type embryos. Five hours after exposure to UVB, embryos lacking gadusol had significantly higher expression of many key stress response genes (*tp53, gadd45aa, ddb2*, & *cdkn1a*) relative to UVB-treated controls (**Fig. 2D**). GO terms enriched in UV-exposed gadusol-depleted embryos included response to UV, response to DNA damage, response to light, and other stress response terms (**Fig. S8, Table S1**). Together, our imaging and gene expression data confirm that gadusol in zebrafish embryos acts as a true sunscreen to provide efficient protection against UV-induced DNA damage, cellular stress, and cell death.

### Gadusol is the primary sunscreen in early fish development

In light of our finding that gadusol acts as a sunscreen, we compared the relative sunscreening potency of gadusol compared to other potential UV-blocking/absorbing mechanisms in larval zebrafish. Melanin is a well-known sunscreen in many organisms including humans. In zebrafish, melanophores become pigmented around 36 hpf, ultimately forming stripes that partially cover the larval brain and body, a pattern that is stable until ~14 dpf^21,22^. Melanophores protect the hematopoietic niche in larval zebrafish^6^, but their role as a whole-body sunscreen remains untested. The *nacre/mitfa* mutant disrupts a key melanophore master regulator and lacks melanophores. We generated two groups of larvae, each with pigmented and unpigmented siblings. One group contained no maternal gadusol, while the other group contained gadusol (**Fig. 3A**). We treated all 5 dpf larvae with 2.5 kJ of UVB and assessed survival in the nursery at 28 dpf. Larvae with gadusol were highly resistant to UVB stress, regardless of pigmentation status (**Fig. 3B**). All larvae that lacked gadusol were highly sensitive to UVB, and larvae that lacked both gadusol and melanin were slightly more sensitive to UVB than their pigmented siblings. At a lower UVB dose (1.5 KJ), we also found a modest but significant effect of melanophores in protecting against UVB (**Fig. S9**). We conclude that while melanin plays a minor role in UVR protection, gadusol is the primary sunscreen in early fish development.

**Figure 3.**
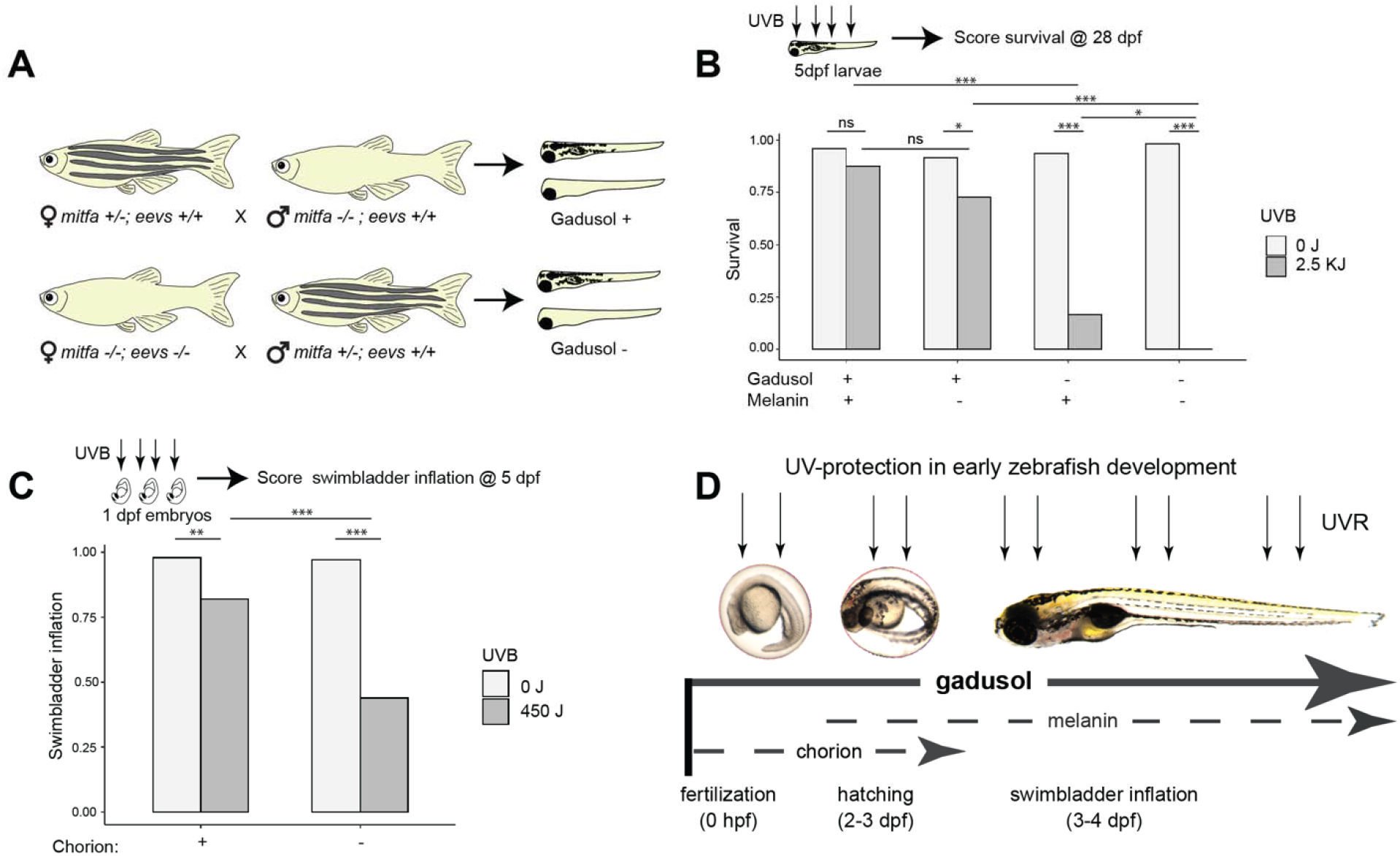
Melanin and the chorion serve as secondary UV-shielding mechanisms in embryonic and larval fish. **A.** Experimental diagram for generating embryos that lack either melanin, maternally provided gadusol, or both. **B.** Survival distribution scored at 28 dpf, with presence of melanin and gadusol indicated, after mock exposure (grey) or UVB exposure (dark grey) at 5 dpf. From left to right, n = 48, 48, 48, 48, 48, 36, 60, 36; N = 2 for each group. **C.** Distribution of swimbladder inflation scored in 5 dpf larvae after mock exposure (grey) or UVB exposure (dark grey) at 24 hpf stage, with or without chorions. n = 100 for each group. N = 3 for each group. **D.** Model illustrating the relative importance and timing of multiple UV-shielding mechanisms used in early zebrafish development. Student’s T-test *p<0.05 ** p<0.01 ***p<0.0001. n = total number of individual embryos/larvae. N = total number of clutches.

Another potential UV-protective mechanism is the chorion, the nearly transparent eggshell that contains perivitelline fluid and the embryo from fertilization until 2-3 dpf. We tested the sunscreening role of the chorion by mechanically removing it with forceps and exposing these embryos, and sibling controls that retained the chorion, to 450 J of UVB at 24 hpf. We found that the chorion does provides significant protection from UVB as ~60% of dechorionated embryos failed to inflate their swim bladders, significantly less than sibling controls (**Fig. 3C**). We examined if gadusol was present in the chorion or in the fluid within the chorion but found little to none (**Fig. S10**). These results suggest that the chorion structure itself can shield some incoming UVB. However, we conclude that the chorion provides less UV protection than gadusol, as gadusol-depleted embryos – even with intact chorions - all failed to inflate their swim bladders when challenged with the same dose of UVB (**Fig. 1C**).

Together, our findings support a model where embryonic and larval fish are protected by multiple layers of UVB protection that span early development (**Fig. 3D**). The egg is maternally loaded with gadusol, which provides the primary and most important layer of UV protection from fertilization until at least 5 dpf. The chorion and melanophores are secondary, and less effective, means of UVR protection. The chorion protects the developing embryo between fertilization and hatching (2-3 dpf), when pigmented melanophores emerge and modestly protect the growing larval fish.

### Gadusol has been repeatedly lost in fish species whose embryos are no longer exposed to sunlight

The two-enzyme biosynthetic pathway necessary for gadusol production (Eevs and MT-Ox) is encoded in numerous vertebrate genomes, including fish, birds, reptiles, and amphibians^17^. Osborn et al. identified the loss of the gadusol biosynthetic pathway in the coelacanth genome, and suggested the loss might be attributable to lack of UV penetration in the deep sea habitat of this species^17^. To test for broader patterns of conservation and loss among fish, we surveyed additional genomes, including many species that live in habitats not exposed to UVR. We hypothesized that gadusol pathway genes would not be required in species that live in deep waters, caves, are live bearers, or use electroreception to navigate habitats with poor light penetrance^23^. To test this hypothesis, we searched 136 teleost genomes for inactivation or loss of either *eevs* or MT-Ox. In all species, we identified a syntenic genomic region demarcated by highly conserved flanking genes and assessed the presence or absence of intact ORFs encoding functional copies of *eevs* and MT-Ox. Our approach largely confirmed that the vast majority of teleosts have functional copies of *eevs* and MT-Ox^17^. However, our survey identified 16 independent losses of either the *eevs* or MT-Ox genes across the teleost phylogeny (**Fig. 4**, red species). Most of these genomes had lost orthologs of both *eevs* and MT-Ox, while others had lost only one gene or had pseudogene remnants. The loss of genes involved in gadusol production was significantly correlated with lifestyle traits that identified species that live or spawn in habitats protected from the sun (p = 0.012) (see **Fig. S12**, **Methods** and **Table S2** for details). To corroborate the link between loss of *eevs* or MT-Ox and loss of gadusol, we measured gadusol levels in medaka embryos, which have intact *eevs* and MT-Ox genes, and ovaries of channel catfish, which have lost *eevs* and MT-Ox. We found a strict correlation between the presence of intact genes and maternally provided gadusol (**Fig. S11**). We conclude that gadusol production has been repeatedly lost during evolution in teleost species whose lifestyles protect them from UVR.

**Figure 4.**
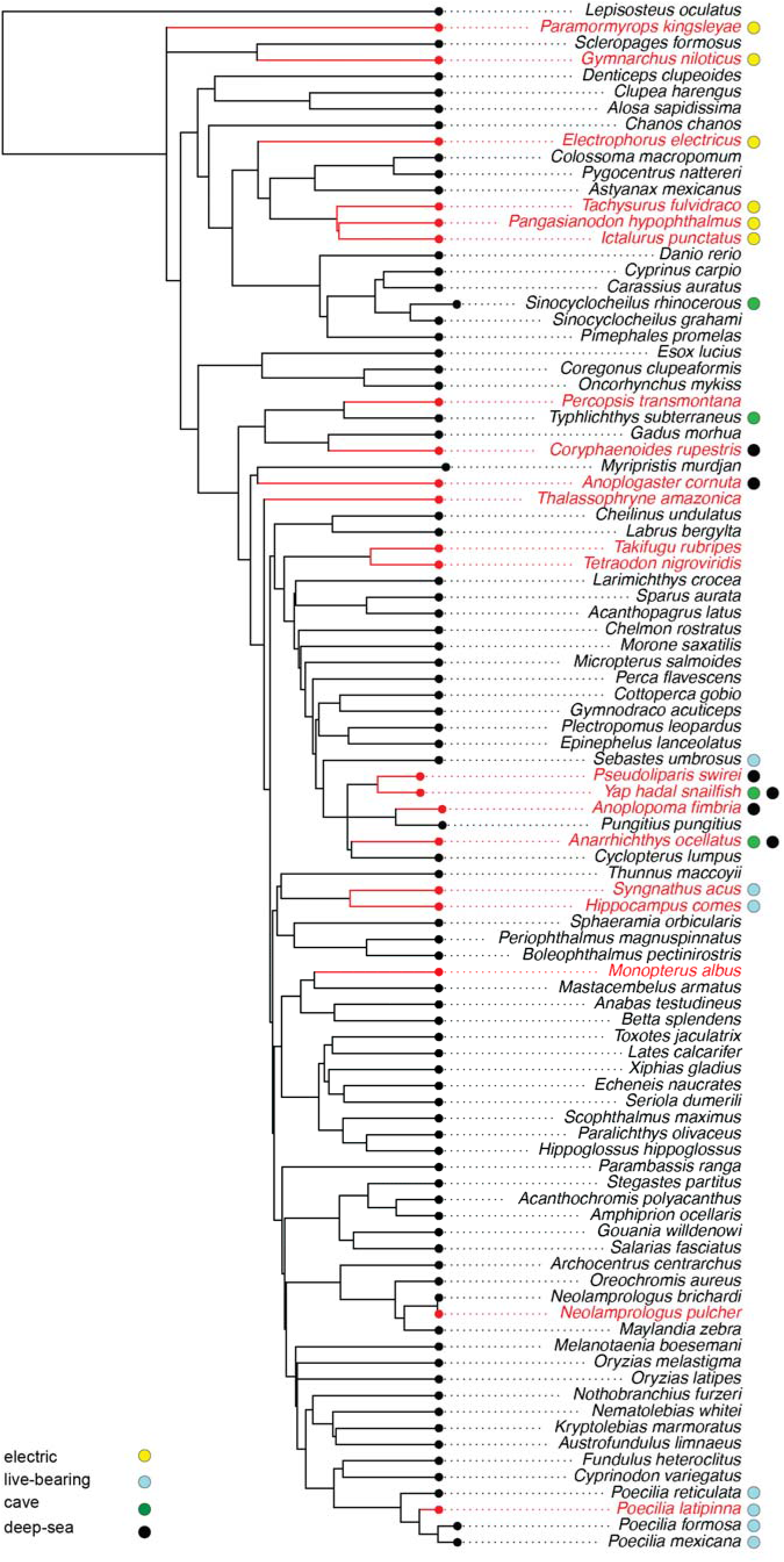
Gadusol production has been lost in several species no longer exposed to UVR. For each of 136 teleost species (full tree in **Fig. S12**), we assessed various life history traits that identify habitats that may not require embryonic protection from UVR, including electroreception, live-bearing, cave dwelling, and deep-sea dwelling, indicated with colors in the legend to the left of the phylogeny. For each species, we identified the presence of intact open reading frames for *eevs* and/or MT-Ox. Species that have lost the genes required for gadusol production are indicated in red. We found 16 independent losses across this phylogeny. We found that fish with these traits are more likely than by chance to lose gadusol (p=0.012).

## Discussion

Plants and microorganisms use numerous UV-absorbing compounds as sunscreens^4,24^. However, other than melanin, the repertoire of vertebrate sunscreens – especially compounds that protect the most vulnerable early stages of development – remain essentially unknown. Here, we provide experimental and phylogenomic evidence that gadusol is an ancient sunscreen essential for protecting fish embryos from UVR. First, we use a CRISPR mutant that disrupts gadusol biosynthesis to show that gadusol is produced during oogenesis and persists in the embryo until at least 5 dpf. Second, we demonstrate that maternally deposited gadusol safeguards embryonic and larval development by preventing UV-induced developmental defects and improving survival. Third, we find that gadusol acts as a true sunscreen preventing the formation of CPDs, a signature of UVB-induced DNA damage, and consequently reducing levels of cell and organismal death. Gadusol does not have any obvious functions beyond protecting against UVR, as mutants survive to adulthood and are fertile. Together, these data demonstrate that gadusol is a maternally provided sunscreen employed during early fish development.

Our work explores two alternative mechanisms of UV protection during early development. We find that the chorion, a transparent eggshell that shields the developing embryo, also provides modest UV protection during embryogenesis. This protection is short lived (zebrafish hatch by 2-3 dpf) but may provide secondary protection during the most vulnerable stages of development. Melanin pigmentation emerges around embryo hatching and serves a relatively modest role as a whole-body sunscreen in 5 dpf larvae. Together, our results show that gadusol is the primary sunscreen across embryonic and larval development, while melanin and the chorion play secondary roles during distinct phases of development.

Finally, our phylogenetic analysis of gadusol biosynthetic genes, building on a previous study^17^, suggest that gadusol is an ancient sunscreen conserved broadly to protect teleost embryos. However, gadusol production has been repeatedly lost during teleost evolution. Intriguingly, these genes are absent in many fish species whose embryos are not exposed to UVR, including deep sea-dwelling and electroreceptive fish. We suggest that similar to our protected fish facility environment, gadusol is also dispensable for embryonic development in natural environments that lack UVR. In microorganisms, the production of sunscreening compounds have been estimated to require >10% of all metabolic activity^24^. Perhaps the loss of gadusol production in nutrient-poor dark habitats provides some evolutionary advantage, analogous to the energy conservation hypothesis invoked to explain the repeated loss of eyes in Mexican cavefish^26,27^. However, once these genes have been lost, descendent species may enter an evolutionary fitness trap where they are confined to breeding environments lacking UVR.

It remains unclear what role gadusol might play in other tetrapods. Functional copies of *eevs* and MT-Ox have been found in numerous vertebrate genomes^17^, but to our knowledge the presence of gadusol has never been reported in vertebrates other than fish. Gadusol has been detected in the eggs or embryos of several aquatic invertebrates, including sponge^28^, starfish^28^, sea urchin^29^, and brine shrimp^30^. We hypothesize that gadusol may also protect early development in these diverse aquatic organisms.

Here, we show that aquatic vertebrates produce and employ an additional sunscreen to melanin. Melanin and gadusol both absorb well in the UVB spectrum. However, melanin also absorbs most wavelengths in the visible light spectrum, making it opaque and conspicuous while gadusol is transparent and invisible. Transparency as camouflage is a common trait in aquatic animals, especially in the open ocean where there is nothing to hide behind^31^. To date, gadusol has only been detected in aquatic organisms. We speculate that gadusol has been particularly advantageous to these animals as it offers protection from UVR, enabling an organism to stay in nutrient-rich sunlit areas, while remaining optically inconspicuous. We propose that aquatic ecosystems exhibit unique ecological challenges that have selected for the use of a transparent sunscreen.

## Materials and Methods

### Zebrafish Husbandry

All zebrafish work was performed at University of Utah’s CBRZ zebrafish facility. This study was conducted under the approval of the Office of Institutional Animal Care and Use Committee (IACUC no. 18-2008) of the University of Utah’s animal care and use program.

### Generation of eevs mutant line

To generate a stable gadusol-depleted mutant line, *eevs* was targeted using CRISPR-Cas9 mutagenesis. Four gRNAs (**Table S3**) were designed using ChopChopV2^32^, targeting exon 2 (**Fig. S1**) due to the lack of suitable target sites within the small exon 1. Guide RNAs were synthesized from DNA oligos using standard protocols^33^. Freshly laid wild-type TU-strain embryos were injected with SpCas9 protein (NEB) mixed with gRNAs (~300 ng/ul), KCl, and phenol red. 1-2 nanoliters were injected into each embryo. Mosaic mutant embryos were raised to adulthood and outcrossed to wild-type Tübingen strain. Primers designed from ChopchopV2^32^ were used to amplify the region targeted for CRISPR editing and to select for edited alleles with large deletions. A compound deletion allele was identified by Sanger sequencing that removes 379bp and shifts the *eevs* open reading frame (**Fig. S1**, sequences in **Table S3**) (Genewiz). This *eevs* mutant allele was given the designation zj2 and can be genotyped using PCR with allele specific primers (**Table S3**). Sibling fish with the zj2 allele were crossed to produce homozygous KO *eevs*^zj2/zj2^ fish, labeled as *eevs*^-/-^ throughout the text. Lack of gadusol was determined using UPLC MS/MS and spectrophotometry (see below for details). Embryos resulting from crosses of *eevs*^-/-^ mothers had little to no gadusol compared to wild-type embryos, confirming successful generation of a gadusol-depleted line.

### Gadusol extraction and UPLC MS/MS detection

Gadusol was extracted twice from embryos (7.5mg of vacuum dried egg material, crushed with a microfuge pestle) using 150 ul of a (80:20, v/v) methanol:water solution. The extraction supernatant was analyzed using ultraperformance liquid chromatography (Waters Acquity I-Class, 2.1 x 100 mm BEH Amide column) and mass spectrometry (Waters Xevo G2 QToF) (UPLC-MS) in negative ionization mode (detector range of 50-2000 Da). We used a regular phase chromatography method starting with 95% acetonitrile (+0.1% formic acid) and 5 % water (+0.1% formic acid) following a linear gradient over 12 minutes ending with 30% acetonitrile (+0.1% formic acid). Analytical standards of pure gadusol were run during the same acquisition run to match the retention time and observed mass between embryo samples and the pure standard.

### Gadusol detection via Nanodrop

To monitor gadusol production the UV-vis spectrometry on a Nanodrop was employed to determine relative gadusol concentrations. Briefly, 25 embryos/larvae were placed in a microfuge tube. All excess water was removed with a Pasteur pipette. 100 ul of 80:20 methanol:water was added to embryos. Embryos were mashed with a microfuge pestle for 15 seconds. Samples were left to extract for at least 15 minutes, and then centrifuged at 12,000 g. Clear supernatant, containing polar compounds such as gadusol, was separated and analyzed on the nanodrop.

### UV exposure, swim bladder inflation, and survival assays

24 hpf embryos were exposed to 450 J of UVB as measured on a radiometer (Solarmeter UVB) at a fluence rate of 2.5 W/m^2^ in 30ml of clear E3 media. This is a conservative estimate of a physiologically relevant UVB dose that fish embryos would routinely experience in the wild^6^. A raised and inverted UVP transilluminator with 306 nm broadband UVB bulbs was used (Ushio 30000318) on the “low” setting (see **Fig. S13**). Embryos were returned to the incubator and kept in the dark after mock or UV exposure. Swim bladder inflation was scored at 5 dpf by adding ice to the petri dish to stun the larvae, followed by manual counting on a dissection scope. A standard dose curve was conducted to determine that 450 J was an appropriate dose (**Fig. S3**).

5 dpf larvae were exposed to a dose curve to determine that 2.5 kJ was an appropriate dose (**Fig. S5**). After mock or UV exposure, larvae were placed in an incubator for 1 day (dark) and then placed in the nursery at 6 dpf. Survival was scored at 28 days post-fertilization to ensure that all living juveniles could feed on their own and were not being sustained on maternal yolk.

### Determination of CPDs in 24 hpf embryos

24 hpf embryos were dechorionated to obtain more consistent UV exposure. Embryos were exposed to 450 J of UVB and then immediately fixed after exposure in 4% PFA for 1 hour at 25°C. Fixed embryos were then washed in PBST. Embryos were exposed to 2 M HCl for 1 hour to break apart dsDNA and expose CPD epitopes. Samples were blocked in 5% NGS + PBST. Mouse anti-CPD primary antibody (TDM-2, Cosmo Bio) was used to stain for CPDs. Goat anti-mouse AF546 secondary antibody (Invitrogen) was used to visualize CPDs. Embryos were also stained with DAPI to visualize nuclei. Prior to imaging on a confocal microscope, tails were removed from embryos and placed on a flat glass slide with a small drop of PBST. A cover slip was mounted over the tails and sealed with nail polish. Tails were then imaged on an inverted confocal microscope with a 20x objective (Zeiss 880). Images were analyzed using ImageJ^34^ to determine mean fluorescence intensity / tail area using the DAPI channel to create a mask for the tail.

### Apoptosis assay

24 hpf embryos within chorions were exposed to 450 J of UVB and then placed in the incubator for 5 hours. Chorions were removed and embryos were fixed for 1 hour in 4% PFA. Embryos were stained with an activated caspase-3 antibody (BD Biosciences, anti:Rabbit) to mark apoptotic cells. Goat anti-rabbit AF594 secondary antibody (Invitrogen) was used to visualize apoptotic cells. Embryo tails were removed, processed, and imaged as above. ImageJ was used to process images and count the number of activated caspase-3 positive nuclei/mm^2^.

### RNAseq sample prep, library prep, sequencing, and analysis

After 5 or 24hrs post UV exposure embryos were smashed with a microfuge pestle (MTC Bio) and RNA extracted using TRI Reagent (Zymo) and purified via Direct-zol RNA Miniprep Plus (Zymo). Library prepared using NEBNext Ultra II Directional RNA Library Prep with poly(A) mRNA Isolation. Samples then sequenced with Total RNA (eukaryote) NovaSeq SP Reagent Kit v1.5_50×50 bp. Each sample sequenced to a depth of 25 million reads. Reads aligned using STAR^35^ and zebrafish reference genome (GRCz11). Optical duplicates removed and adapters trimmed. Differential expression analysis conducted with DESeq2^36^ and specifically the Bioconductor package^37^.

### Generating embryos that lack melanin and gadusol

To generate embryos that lacked melanin, *mitfa^w2/w2^* fish were crossed with *mitfa^+/w2^* fish to produce clutches of 1:1 pigmented:unpigmented siblings, all with gadusol (**Fig. 3A**). To generate embryos that lack both melanin and gadusol, *mitfa^+/w2^; eevs*^-/-^ females were crossed to *mitfa^w2/w2^; eevs*^-/-^ males to produce 1:1 pigmented:unpigmented siblings that all lacked gadusol.

### Chorion UV protection assay

24 hpf wild-type TU-strain embryos were manually dechorionated with forceps in a dish with a thin film of 0.5% agar on the base of the dish. Embryos were moved with a fire-smoothened Pasteur pipette. Embryos were exposed to 450 J of UVB as described above and then placed in incubator and swim bladder inflation was scored at 5 dpf.

### Phylogenetic analysis of eevs and MT-Ox presence

123 genomes were gathered from the UCSC genome ark (GenArk) and additional 11 genomes for deep sea and electro-receptive fish were gathered NCBI genomes for all except the Yap

Hadal snailfish^38^ and *pseudoliparis swirei*^39^. A BLAST database for each species was created by using the zebrafish sequence spanning from FRMD4B to FOXP1 to find the same region in all curated genomes. If there was no BLAST hit for FOXP1 or MITF then the genome was dropped for low quality. We then performed a tBLASTn search on the created databases for the remaining genomes, using the zebrafish EEVS and MtOX translated nucleotide sequence as the query. If there was no hit for EEVS or Mt-OX in the tblastn search that species was labeled as not having gadusol.

To correlate the presence/absence of gadusol with life history traits we first collected life history data for all species (**Table S2**). The life history traits that we annotated were deep-sea, live-bearing, electro-reception, and cave dwelling. We then built a species tree using fishtree^40^ and added the Yap hadal snailfish and *Pseudolapris swirei* using the phylogenetic relationship determined in Mu et. al ^38^. Due to gene loss in sister species not being independent, we used Bayestraits^41^ to perform the correlation test. We used discrete model testing and a likelihood ratios test comparing each of the five life-history traits to loss of gadusol (**Table S2**).

When running Bayestraits the loss of gadusol (parameter beta1 in the independent model and q31 and q42 in dependent model) was set as trait one and the various life history traits were set as trait two. The rate at which gadusol can be regained after loss was constrained to zero because we were scoring for loss of the gene, and assumed it is nearly impossible to regain the gene, especially in the short time span we are investigating. The parameters that estimate the rate of life history traits changing from absent to present (q12,q34,q21,q43) were constrained to equal to each other, under the assumption that it is unreasonable that a fish would change its life-style after loss of gadusol. When comparing the cave life history to gadusol loss, the parameter that estimates the rate of moving from cave to surface (q21 and q43) was constrained to zero under the assumption that species don’t re-emerge from a cave after adapting to that lifestyle.

The significance of the correlation between life-history trait and loss of gadusol was determined using a likelihood ratio test which is calculated by 2*((dependent model likelihood)-(dependent model likelihood)). The significance is then determined using a chi-square distribution with 2 degrees of freedom.

All parameters and code to re-run these models can be found in https://github.com/nclark-lab/gadusol

## Acknowledgements

We thank all members of the Gagnon lab for helpful discussions and comments. We thank Taifo Mahmoud for providing pure gadusol samples, Phyliss Coley for use of her mass spectrometer, Angie Serrano for help with IHC protocols, and Julie Hollien for use of her UVP transilluminator, and Nels Elde, Nitin Phadnis, Alex Schier, and Michael Shapiro for comments on the manuscript. We thank CZAR and CBRZ staff, especially Nathan Baker, for excellent zebrafish care. We thank the Cell Imaging and the HCI Bioinformatics core facilities. This research was conducted on the traditional and ancestral homeland of the Shoshone, Paiute, Goshute, and Ute Tribes. We affirm and support the University of Utah’s partnership with Native Nations and Urban Indian communities.

## Author Contributions

MCR and JAG conceived of the study. MCR created *eevs* knockouts. MCR carried out and designed UV experiments. JM and MCR performed RNAseq experiments. MCR analyzed RNAseq data with U of U bioinformatics core. DLF analyzed and ran samples on UPLC MS/MS. JHL explored fish phylogeny and determined gene loss with input from NLC and MCR. MCR and JAG wrote the manuscript with input from all authors.

## Data and materials availability

All data necessary for evaluating the conclusions in this paper are present in the paper or in supplemental figures. RNAseq data are being deposited in GEO.

