## Supplemental Materials for "Gadusol is a maternally provided sunscreen that protects fish embryos from DNA damage"

**Supplemental Materials for Rice et al., BioRxiv**

**Table of Contents**

| **Figure S1** | Description of the zebrafish eevs mutant allele. |
| --- | --- |
| **Figure S2** | Gadusol is only maternally provided in embryos. |
| **Figure S3** | Swim bladder inflation is highly correlated with survival. |
| **Figure S4** | UVB dosage curve on 24 hpf embryos. |
| **Figure S5** | UVB dosage curve on 5 dpf larvae. |
| **Figure S6** | Gadusol does not impact sensitivity to H_2_O_2_. |
| **Figure S7** | Representative images of zebrafish embryo tails stained for a marker of apoptosis. |
| **Figure S8** | Clusters of differentially-expressed genes emerge in UV-treated embryos that lack gadusol. |
| **Figure S9** | A modest role for melanin as a sunscreen in larval fish. |
| **Figure S10** | Gadusol is absent from the chorion and perivitelline fluid. |
| **Figure S11** | Gadusol is absent in catfish roe and present in medaka embryos. |
| **Figure S12** | Gadusol production has been lost in several species no longer exposed to UVR. |
| **Figure S13** | Experimental setup for UVB exposure. |
| **Table S1** | GO term enrichment from RNAseq. |
| **Table S2** | Life history trait scoring and statistical analyses for gadusol loss across fish species. |
| **Table S3** | DNA sequences for *eevs* mutant. |

**Tables are attached as separate files.**

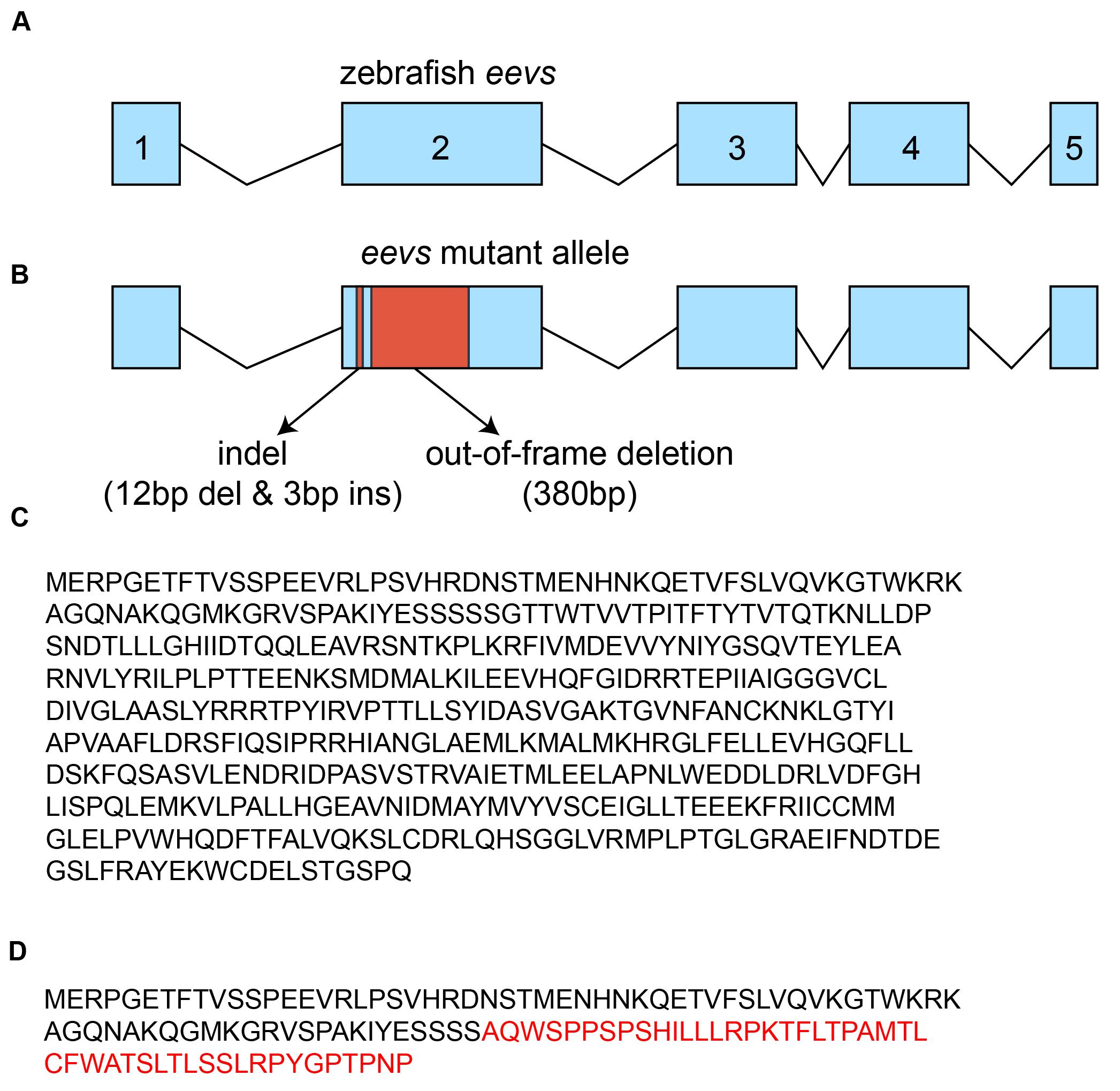

**Figure S1. Description of the zebrafish *eevs* mutant allele.**

**(A)** Diagram of the gene structure of the zebrafish wildtype *eevs* allele, with exons depicted as boxes and numbered and introns depicted as connecting lines. **(B)** Diagram of the mutant *eevs* allele (zj2) containing a 379 bp deletion (red) generated using CRISPR-Cas9, which frameshifts the *eevs* gene. **(C)** Predicted amino acid sequence of wildtype zebrafish *eevs* protein. **(D)** Predicted amino acid sequence of zj2 mutant zebrafish *eevs* protein. The zj2 deletion is predicted to cause a frameshift producing out-of-frame amino acids (red) before a premature stop codon.

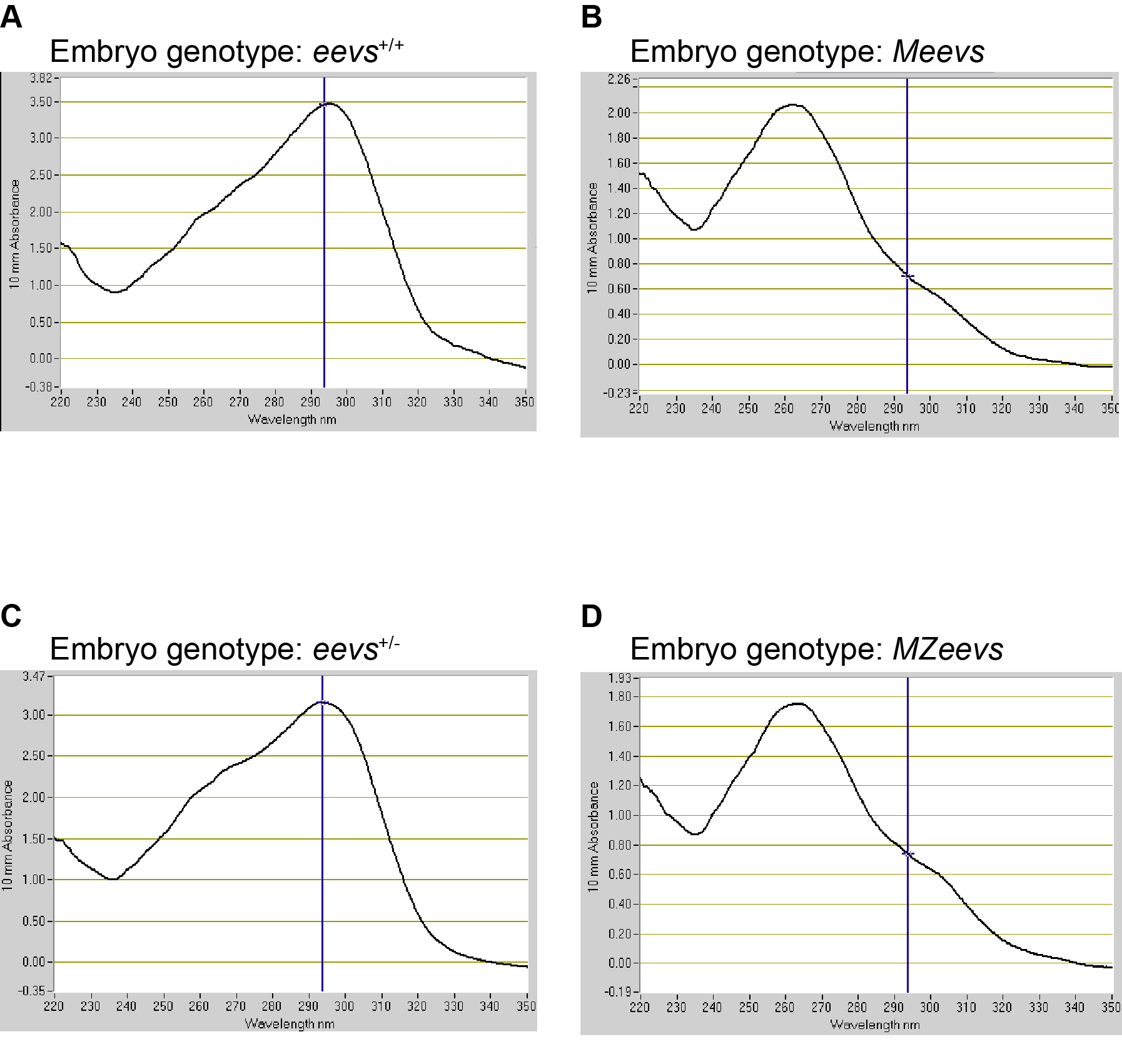

**Figure S2. Gadusol is only maternally provided in embryos.**

Nanodrop UV-spectrograms of polar compounds from methanol-extracted embryos with the indicated genotypes – *eevs*^+/+^ (**A**), M*eevs* (**B**), *eevs*^+/-^ (**C**), and MZ*eevs* (**D**). The absorption peak for gadusol at neutral pH is 296 nm, indicated with a blue line. Absorption at 296 nm is absent in M*eevs* and MZ*eevs* embryos.

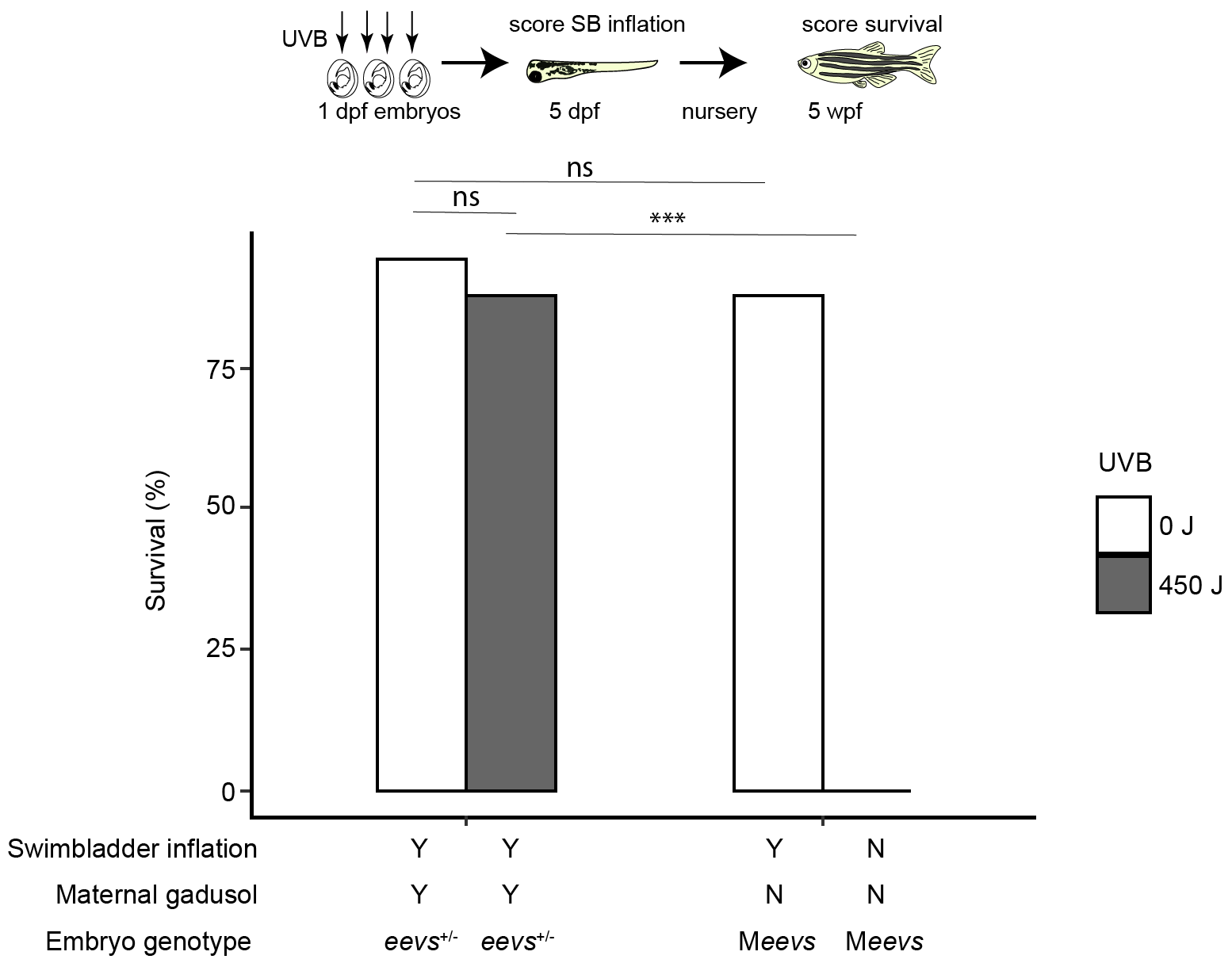

**Figure S3. Swim bladder inflation is highly correlated with survival.**

Diagram of the experimental design. *eevs*^+/-^ and M*eevs* embryos were mock-exposed (white) or exposed to 450 J of UVB (grey) at 24 hpf, scored for swim bladder inflation at 5 dpf, placed in the fish facility nursery, and scored again for survival at 5 weeks of age. Survival rates after mock- or UVB-exposure are shown in the bar graph. M*eevs* embryos exposed to UVB failed to inflate their swim bladders and fail to survive in the nursery, while *eevs*^+/-^ embryos exposed to UVB inflated their swim bladders and survived in the nursery (Fisher’s exact ***p<.0001). Mock-exposed embryos inflated their swim bladders and survived in the nursery at similar rates regardless of genetic condition. From left to right, n = 50, 50 50, 43. Two clutches of embryos were used for each group.

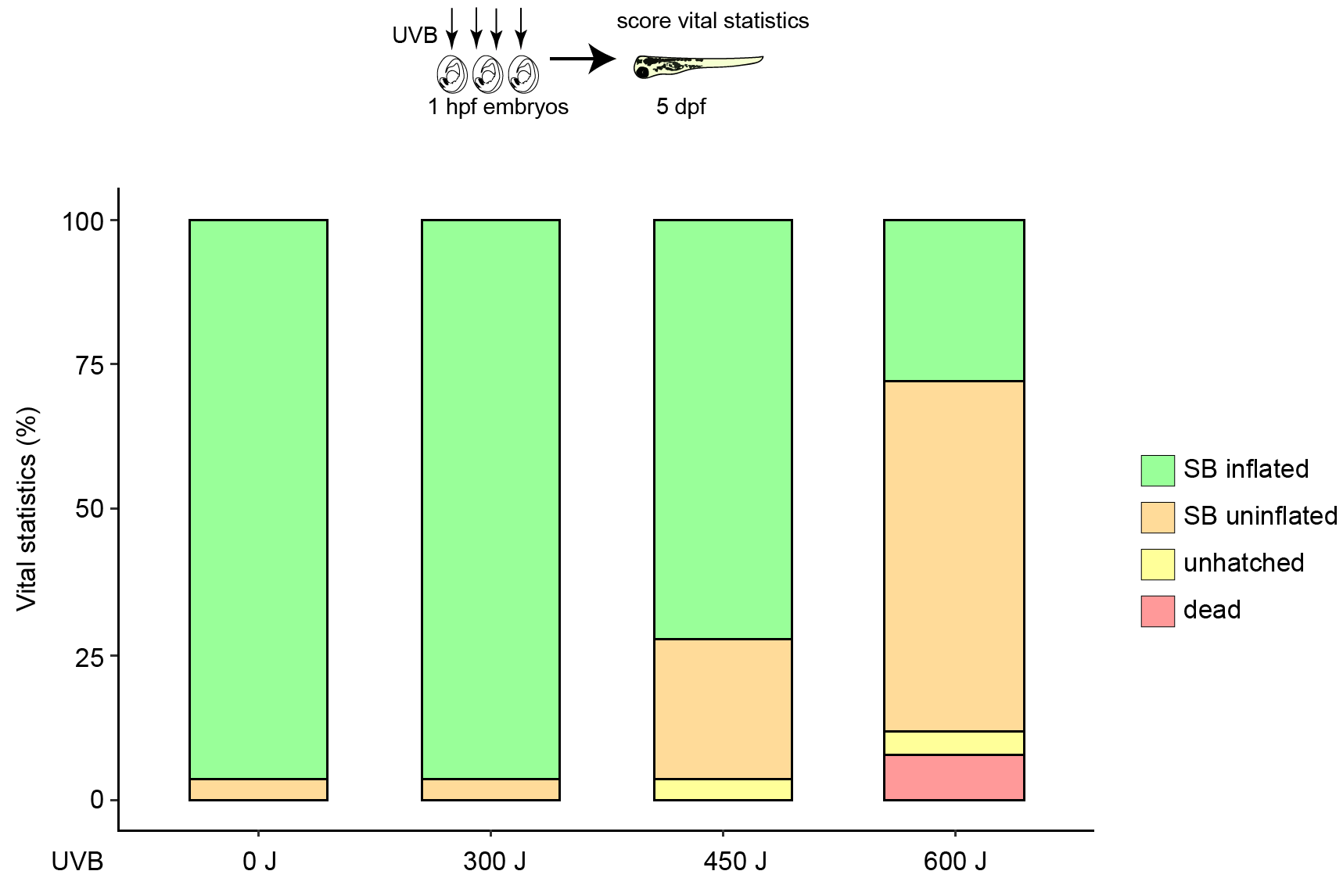

**Figure S4. UVB dosage curve on 24 hpf embryos.**

24 hpf *mitfa*^+/-^ embryos were exposed to the indicated doses of UVB and vital statistics were scored at 5 dpf. We scored swim bladder (SB) inflation, chorion hatching, and obvious mortality. 5 dpf larvae that do not hatch or inflate their swim bladder will not survive (also see **Figure S4**). We generated TU/AB hybrid strain embryos to mimic the background genetics of the M*eevs* and *eevs*^+/-^ embryos used elsewhere in this paper. The parental cross was a wild-type TU strain female to a *mitfa*^-/-^ AB strain male. From left to right, n = 25, 26, 25, 25. Based on these data, the 450 J dose was selected for all UVB exposures to 24 hpf embryos as it resulted in modest defects in swim bladder inflation but did not result in immediate mortality.

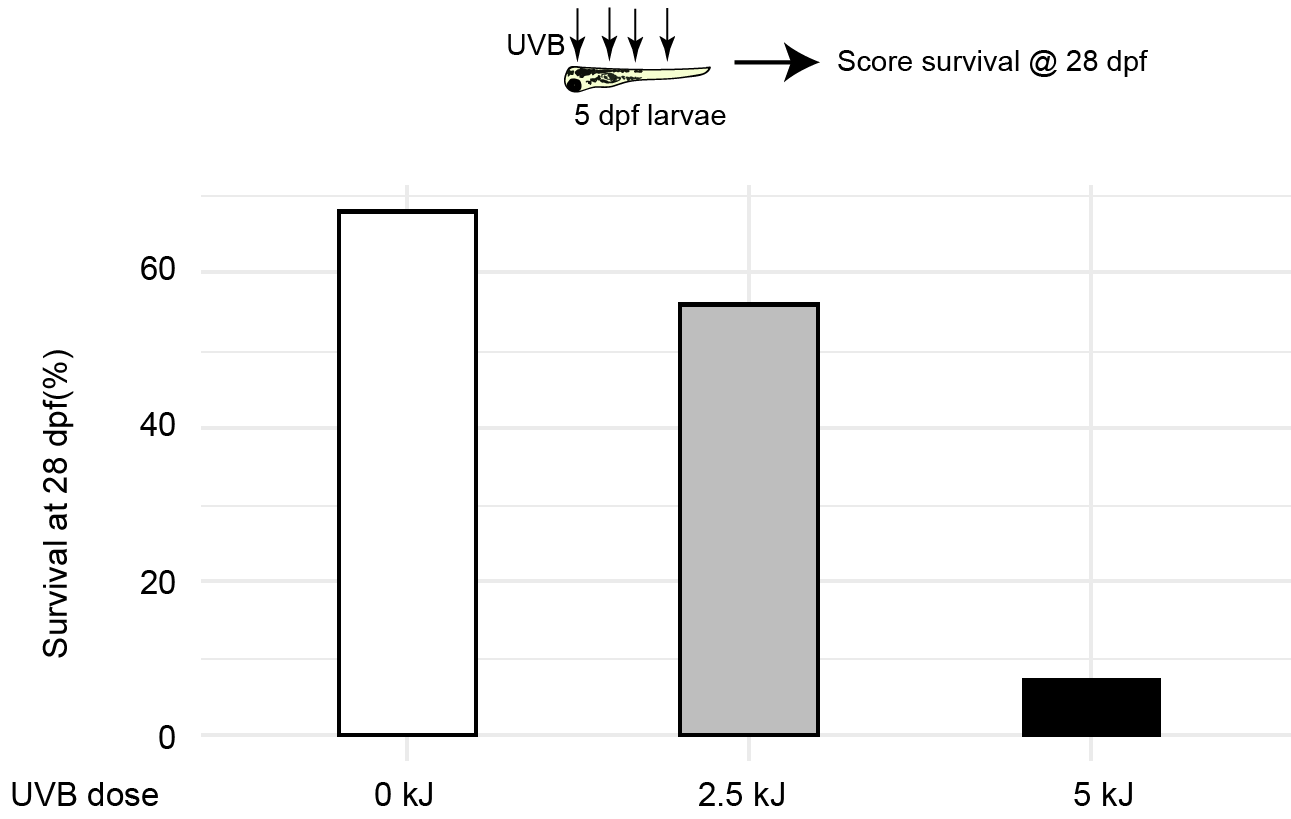

**Figure S5. UVB dosage curve on 5 dpf larvae.**

5 dpf wild-type TU embryos were exposed to the indicated doses of UVB and placed into
the fish facility nursery. Survival was scored at 28 dpf. From left to right, n = 100, 100, 100. Based on these data, the 2.5 kJ dose was selected for all UVB exposures to 5 dpf embryos as it resulted in a modest decrease in nursery survival.

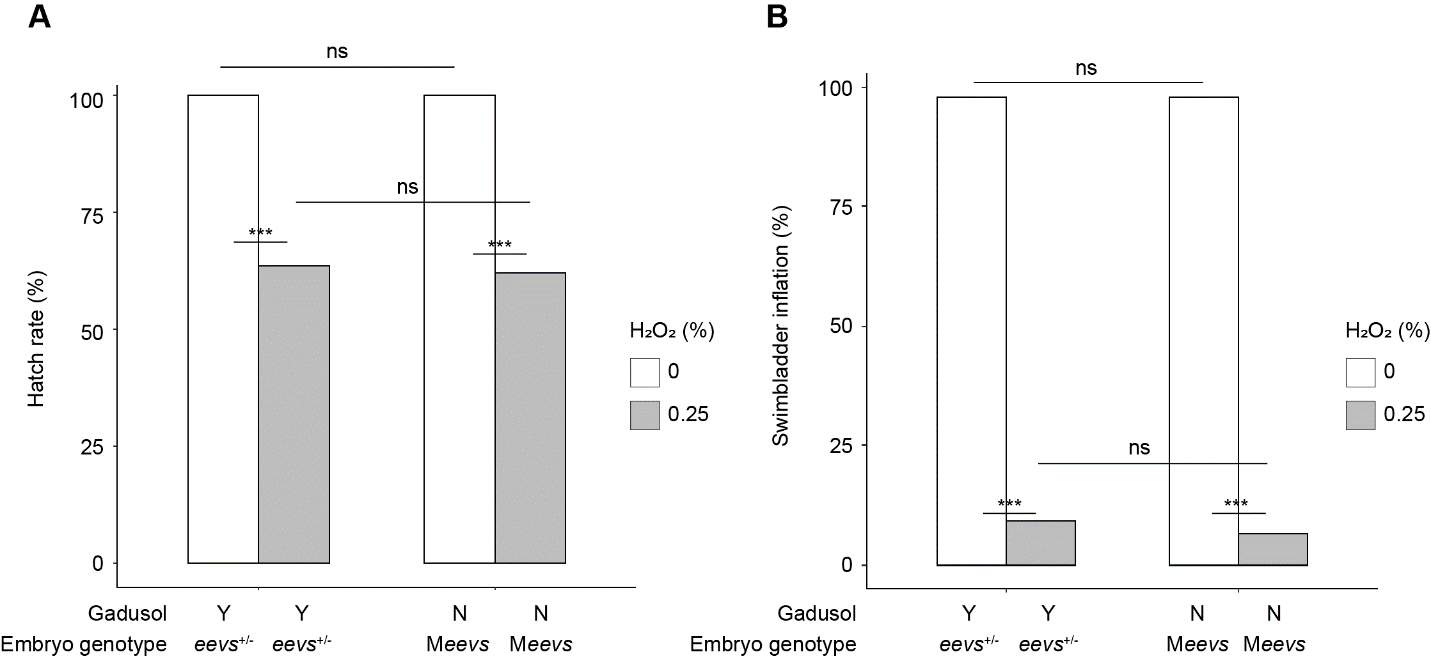

**Figure S6. Gadusol does not impact sensitivity to H_2_O_2_.**

24 hpf *eevs*^+/-^ or M*eevs* embryos were incubated in 0% or 0.25% hydrogen peroxide for 1 hour, then washed in E3 embryo medium. (**A**) Hatch rate displayed between groups showing no difference between embryos with or without gadusol, before or after hydrogen peroxide treatment. (**B**) Swim bladder inflation rate shown. No difference is observed between embryos with or without gadusol, before or after hydrogen peroxide treatment. Hatch rate and swimbladder inflation scored at 5 dpf. Embryos that do not hatch never inflate their swim bladder. All embryos result from TU strain eevs^-/-^ males or females outcrossed with AB strain males or females. From left to right, n = 45, 44, 45, 45. Two clutches of embryos were used for each group.

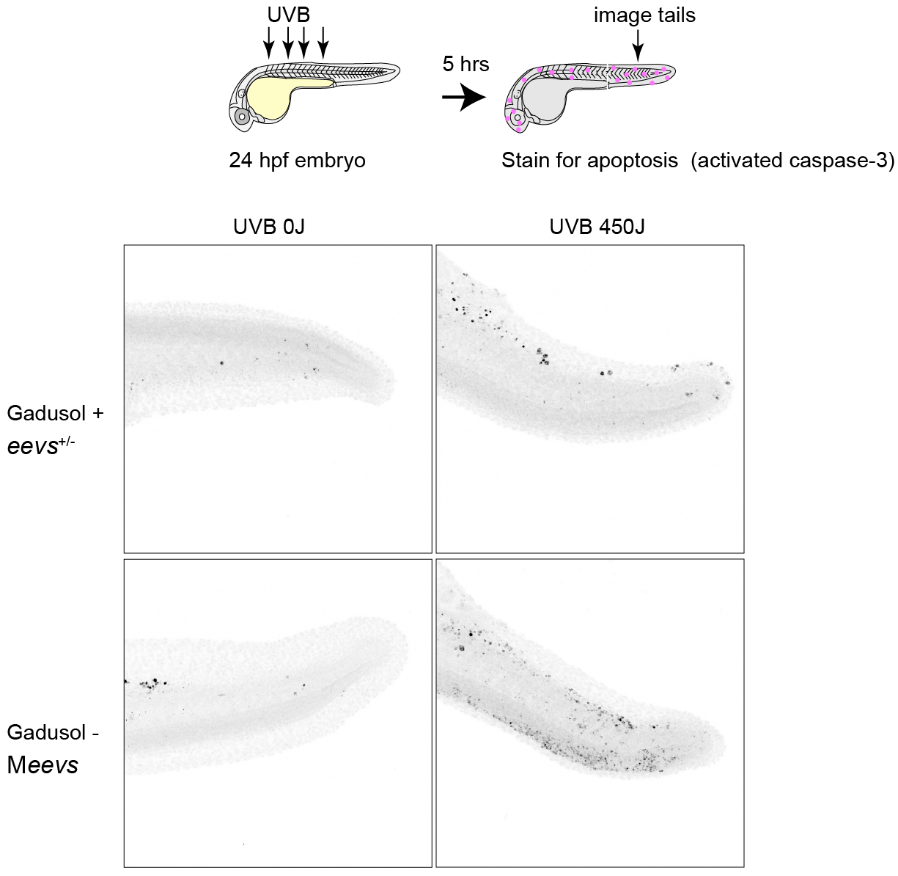

**Figure S7. Representative images of zebrafish embryo tails stained for a marker of apoptosis.**

24 hpf *eevs*^+/-^ or M*eevs* embryos within chorions were mock-exposed or exposed to 450 J of UVB. 5 hours later, embryos were fixed and stained using an activated caspase-3 antibody. Fluorescence intensities were quantified as described in the **Methods** and displayed in **Figure 2C**.

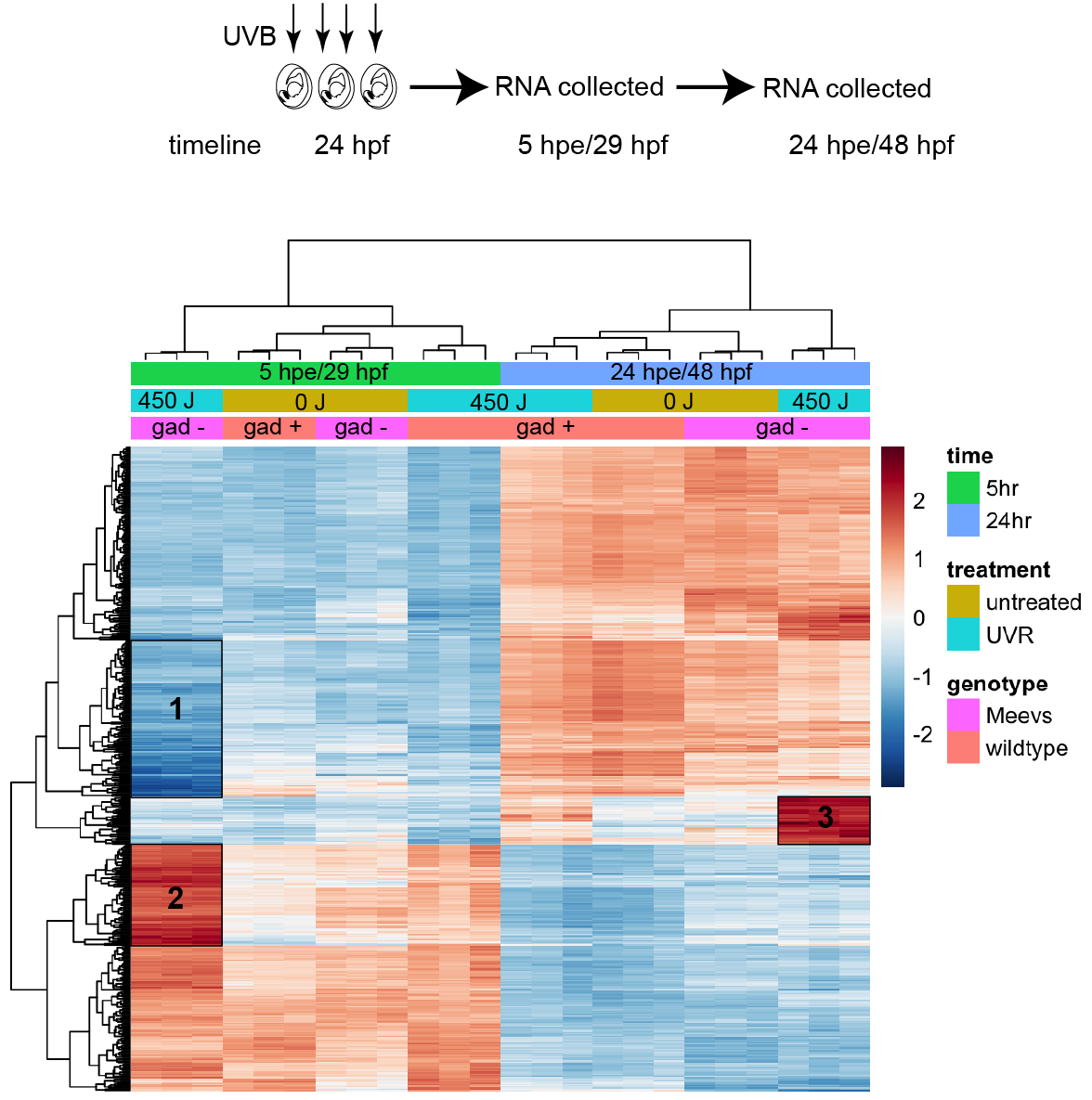

**Figure S8. Clusters of differentially-expressed genes emerge in UV-treated embryos that lack gadusol.**

Displayed is a heat map of the 500 genes with most variable expression levels across conditions. Each row is a gene, each column is a single RNAseq experiment. Each condition (M*eevs* or wild type, mock-exposed or exposed to UVB, 5 hours or 24 hour after exposure) is represented by three replicate RNAseq experiments. Three clusters of genes are highlighted that emerged from unbiased hierarchical clustering. Cluster 1 genes were strongly downregulated in embryos that lack gadusol 5 hours after exposure to UVB. Genes in cluster 1 are associated with GO terms that indicate downregulation of transcription. Cluster 2 genes were strongly upregulated in embryos that lack gadusol 5 hours after exposure to UVB. Genes in cluster 2 are associated with stress response and DNA damage response GO terms. Select genes from cluster 2 are shown in **Figure 2D**. Cluster 3 genes were strongly upregulated in embryos that lack gadusol 24 hours after exposure to UVB. Cluster 3 contains two genes associated with the wound healing GO term, suggesting that the UV-treated gadusol-lacking embryos are still regulating a response to wound healing 24 hours after UV exposure. See SI Table 1 for all GO terms associated with each cluster.

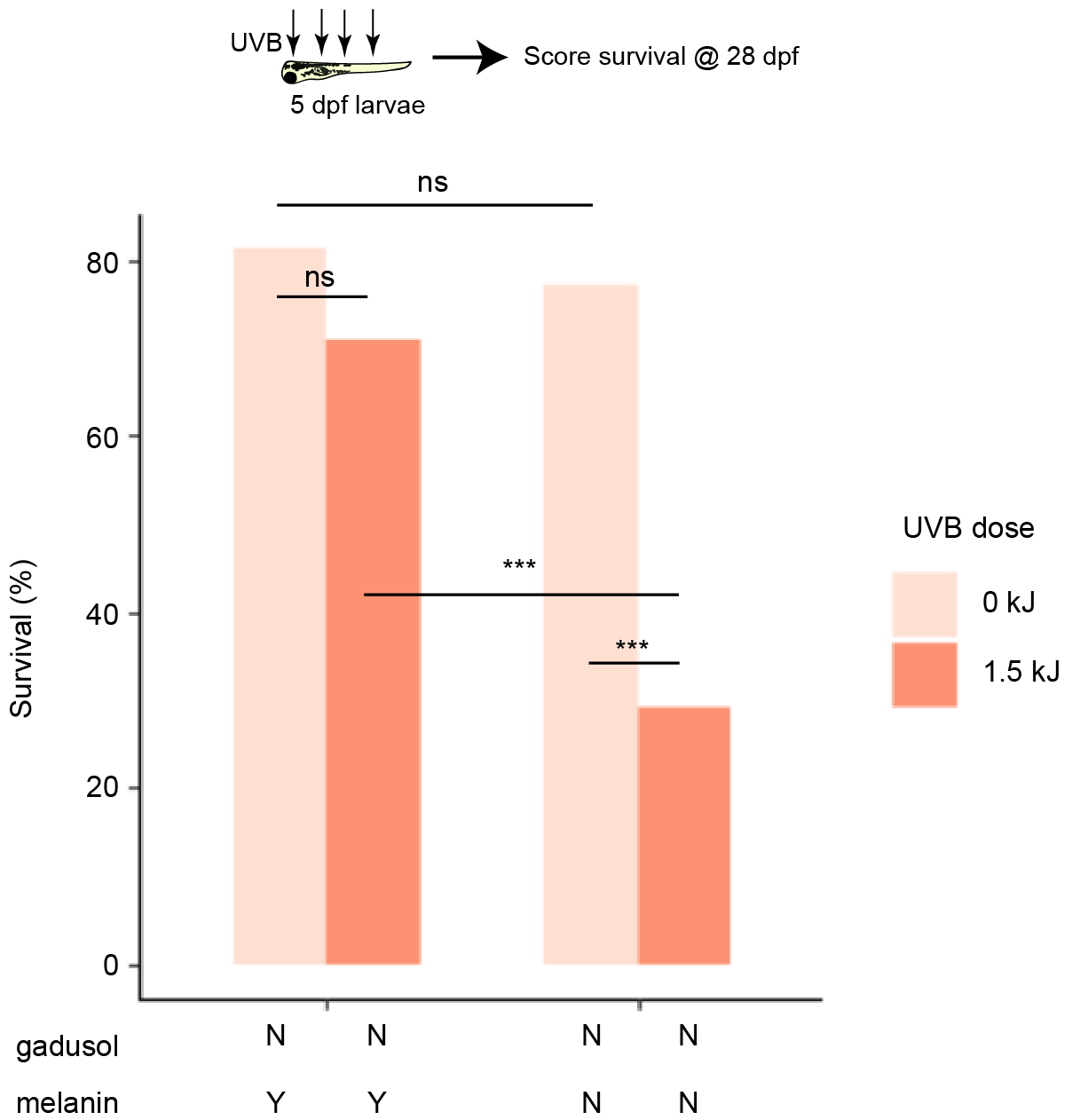

**Figure S9. A modest role for melanin as a sunscreen in larval fish.**

Larvae with or without maternal gadusol, and with or without melanin, were exposed to 1.5 kJ UVB (a lower dose than in other experiments) and placed into the fish facility nursery. Survival was scored at 28 dpf. All larvae are siblings. n = 48 for all groups. Two clutches of embryos were used for each group. Statistics: fisher’s exact *** p<.0001.

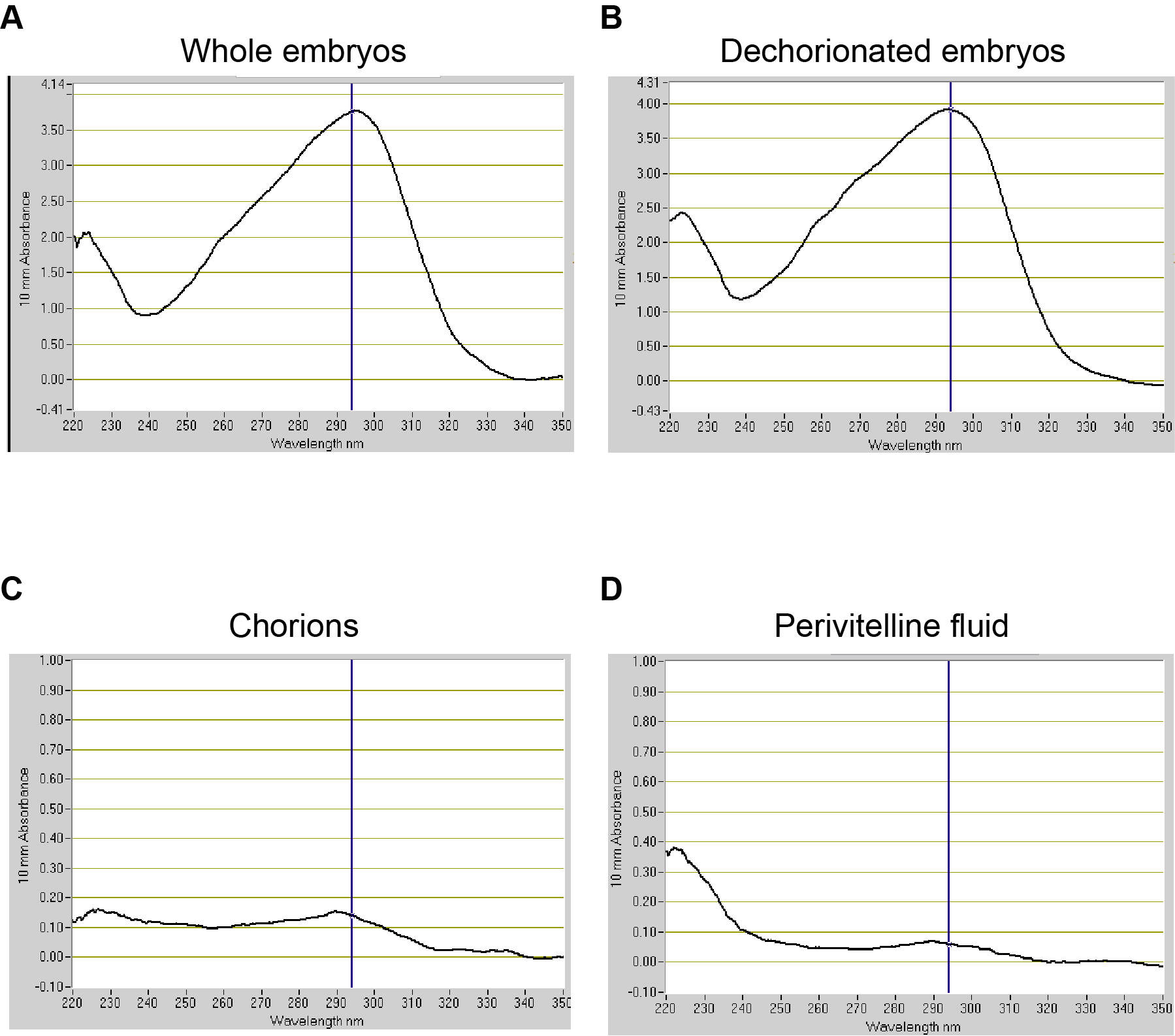

**Figure S10. Gadusol is absent from the chorion and perivitelline fluid.**

Nanodrop UV-spectrograms of polar compounds from the following samples – whole embryos (**A**), dechorionated embryos (**B**), isolated chorions (**C**), isolated perivitelline fluid (**D**). The absorption peak for gadusol at neutral pH is 296 nm, indicated with a blue line. Absorption at 296 nm is absent in chorion and perivitelline fluid samples.

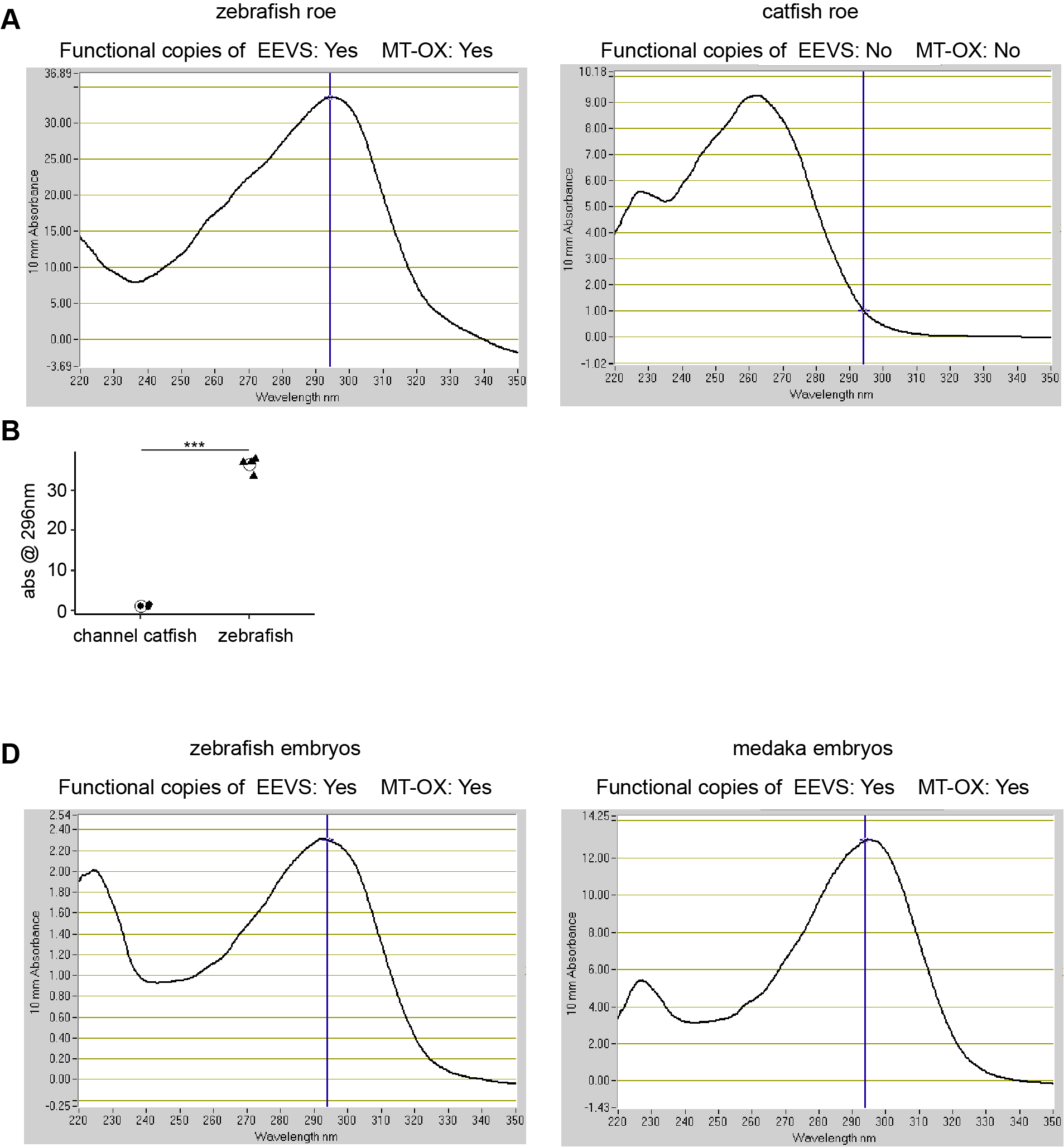

**Figure S11. Gadusol is absent in catfish roe and present in medaka embryos.**

Nanodrop UV-spectrograms of polar compounds from the following samples – Channel catfish (*Ictalurus punctatus*) ovaries (**A**), zebrafish ovaries (**B**), fertilized Medaka (*Oryzias latipes*) eggs (**C**), fertilized zebrafish eggs (**D**). The absorption peak for gadusol at neutral pH is 296 nm, indicated with a blue line. The channel catfish genome does not contain intact copies of *eevs* and MT-Ox, and absorption at 296 nm is absent in catfish ovaries.

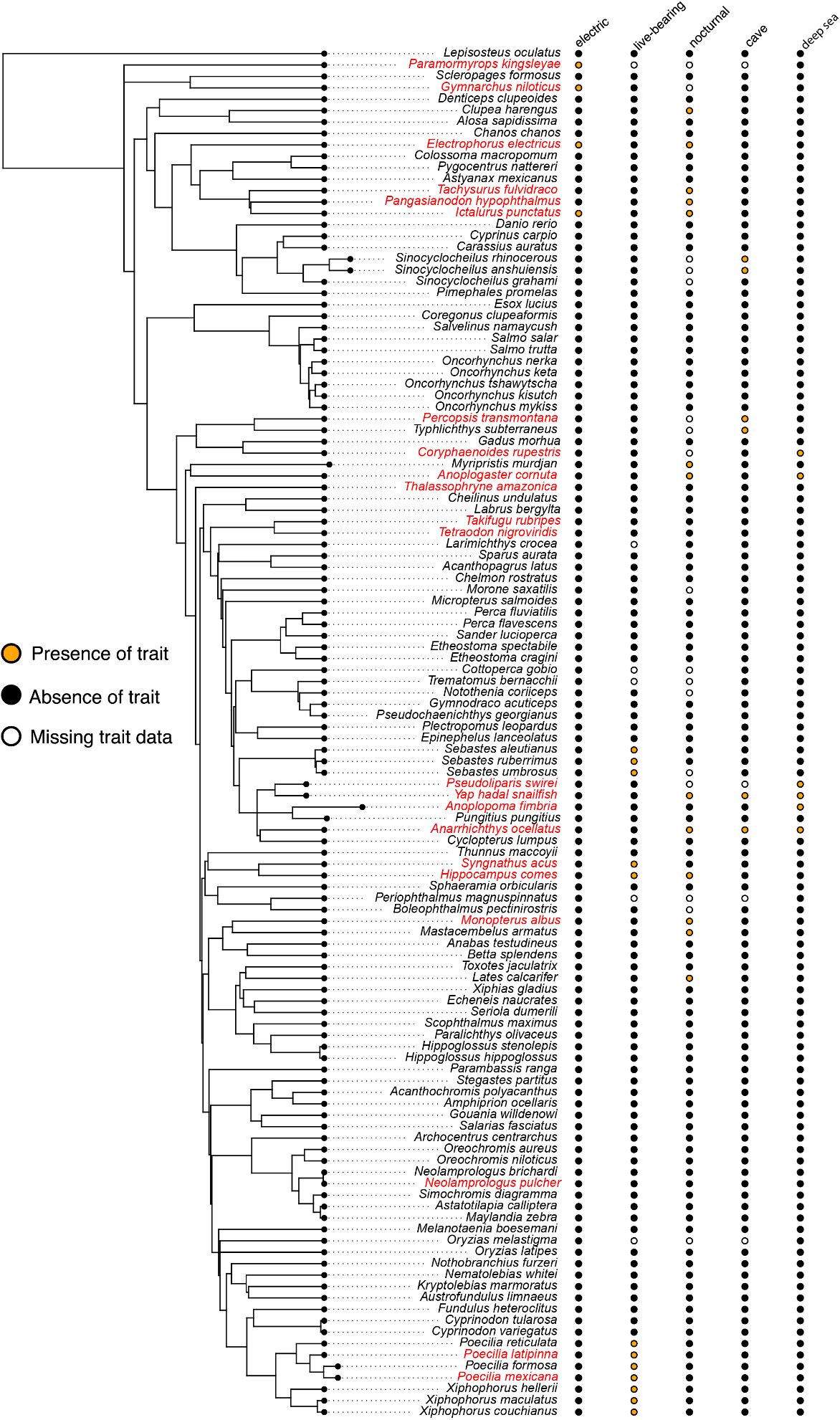

**Figure S12. Gadusol production has been lost in several species no longer exposed to UVR.**

For each of 136 teleost species, we assessed various life history traits that identify habitats that may not require embryonic protection from UVR, including electroreception, live-bearing, cave dwelling, and deep-sea dwelling, indicated with colors to the left of the phylogeny. For each species, we identified the presence of intact open reading frames for *eevs* and/or MT-Ox. Species that have lost the genes required for gadusol production are indicated in red. We found 16 independent losses across this phylogeny. We found that fish with these traits are more likely than by chance to lose gadusol (p=0.012).

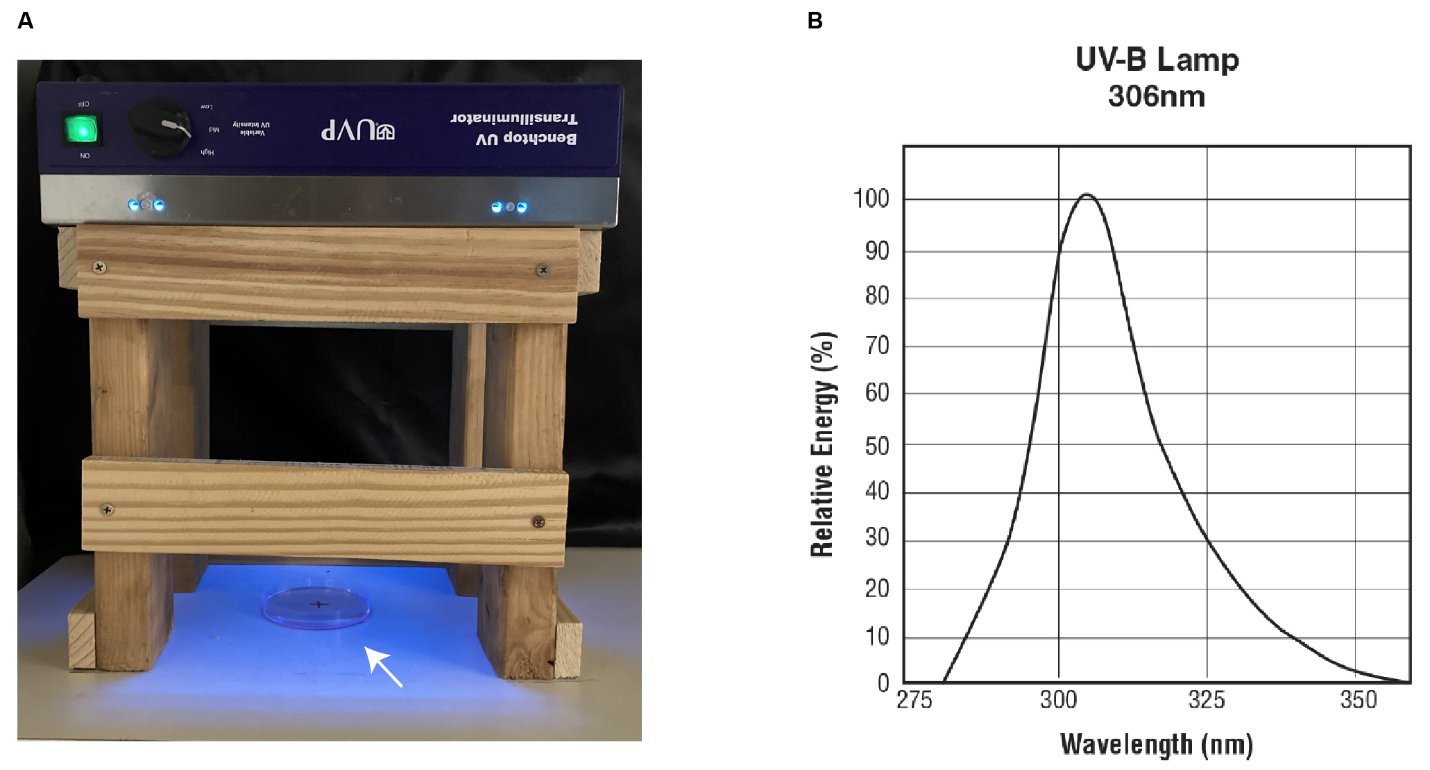

**Figure S13. Experimental setup for UVB exposure.**

**(A)** A transilluminator rests inverted on a 30 cm wooden stand in order to set the fluence rate to 2.5 W/m², an ecologically relevant rate. Arrow denotes location of the petri dish. Embryos and larvae are placed in 30ml of clear E3 buffer. Lids are removed during UV exposure. (**B**) UVB spectrum of 8 W broadband 306nm UVB bulbs (Ushio 30000318). Note absence of UVC light.
